# Plant diversity and water regime have distinct effects on soil fungal communities that are linked to plant productivity

**DOI:** 10.1101/2025.10.10.680786

**Authors:** Xinyi Yan, Elizabeth A. Bowman, Sarah K. Ortiz, Damla Cinoğlu, Caroline E. Farrior, Amelia A. Wolf

## Abstract

The soil microbiome both shapes and is shaped by the ecological consequences of drought and plant diversity loss. Sequencing soil fungal communities in a biodiversity experiment, we found that water addition, plant species richness, and plant phylogenetic diversity all impacted soil fungal composition, and increased the arbuscular mycorrhizal fungi (AMF) richness in particular. The richness of different fungal functional guilds all increased with plant functional trait diversity but were related to different above- and below-ground plant functional traits -- AMF richness was negatively associated with root traits that indicate mycorrhizal dependence, whereas diverse soil pathogens generally co-occurred with plant traits linked to faster life histories. Unexpectedly, the effect of plant diversity loss on fungal diversity did not interact with the effects of either the watering treatment or the community weighted mean of plant traits. Shifts in AMF richness and soil fungal taxa due to plant diversity loss were further associated with reduced plant productivity and diversity effects. Overall, our results indicate that changes in water input and loss in plant taxonomic, phylogenetic, and functional diversity can negatively impact soil fungal communities, and through changes in soil microbiome, may also reduce plant productivity. This negative effect of plant diversity loss is consistent across water regimes and plant community-level traits, suggesting a general vulnerability of soil microbiome and ecosystem productivity towards the biodiversity crisis.

## Introduction

Biodiversity loss and climate change are two major threats to our contemporary ecosystems, and soil microbes could be both the victim and mediator of these threats. Globally, a third or more plant species are under threat of extinction (Nic Lughadha et al. (2020), Pimm and Joppa (2015)). While plant diversity change can be variable in communities (Vellend et al. (2013)), systems facing increased warming and drought often suffer pronounced declines (Harrison (2020), Harrison et al. (2015)), suggesting that continued climate change could further exacerbate biodiversity loss (Bellard et al. (2012)). Both plant diversity loss and drought together can have downstream effects the soil microbial communities, and on ecosystem functions such as productivity. The soil microbiome, on the other hand, may help buffer plant communities against some of these negative impacts (Van Der Heijden et al. (2008), Muhammad et al. (2025)). While increasing studies (Fahey et al. (2023), García-Palacios et al. (2015)) have been tackling the microbial role in mediating ecosystem productivity under global changes, there is still no consensus on this topic.

The first level of complexity lies in potentially different ways soil microbiomes respond to different plant diversity loss scenarios. Plant diversity loss can change local soil microbial diversity, an important indicator of ecosystem function and resilience. However, plant diversity can be measured in multiple ways. Loss of species richness, the simplest yet most inclusive metric, corresponds to the scenario where species are lost randomly; and loss of phylogenetic diversity (PD) captures conserved loss of evolutionary history, especially with species-specific soil pathogens and mutualists (Lu et al. (2024)); loss of functional diversity more directly reflects the loss of mechanism of plant-microbe interactions mediated by plant traits. The latter two metrics have been proposed to be better predictors of ecosystem functioning than species richness (Cadotte et al. (2008)) and may better reflect realistic biodiversity loss, where closely related or functionally similar species tend to be lost together (Willis et al. (2008), Harrison et al. (2015)). However, despite the many observational studies, experiments simultaneously comparing the effect of multiple means of diversity loss remain rare (but see Fahey et al. (2023)).

The second level of complexity lies in the inconsistent effect of plant diversity on soil microbial diversity due to context dependency. Across various plant diversity measures and ecological scales, the relationship between plant diversity and soil microbial richness can be highly variable (Liu et al. (2020), Wardle (2006)), ranging from positive (Schmid et al. (2021), Fahey et al. (2023)), absent (Prober et al. (2015), Tedersoo et al. (2014)), or even negative (Schlatter et al. (2015)). Both abiotic and biotic factors likely contribute to these variable relationships. First, water availability can not only affect microbial communities themselves (Schimel (2018), Bogati and Walczak (2022), Malik and Bouskill (2022)), but also mediate the effect of biodiversity loss on soil microbes. For example, increased water availability may strengthen the positive relationship between plant and microbial diversity – under drought, microbial communities may be dominated by fewer taxa with high stress tolerance throughout the plant diversity gradient, whereas higher water availability could reduce environmental filtering, intensify microbial competition, and increase niche differentiation with plant diversity. Second, not only the functional diversity, but also the dominant strategy of plant communities, can affect microbial diversity. Pathogen richness is predicted to increase with “faster” plant strategies such as higher specific leaf area (thinner leaves) and lower root tissue density(Wright et al. (2004)), whereas mycorrhizal fungi richness is predicted to increase with more “outsourcing”, mycorrhizal-dependent strategies of larger root diameter and lower specific root length (Bergmann et al. (2020)), which provide more root space for mycorrhizal colonization. Likewise, plant strategy can also dictate the effect of water availability and plant diversity on microbial diversity. For example, we may expect mycorrhizal richness to increase with the diversity of more “outsourcing” plants that are more likely to foster mycorrhizal partners. Likewise, plant pathogen richness may increase more with the diversity of “faster”, disease-prone plants, compared to that of “slower” plants.

The multidimensional nature of the microbial composition imposes the third layer of complexity in characterizing the microbial response to and effects on plant communities. The changes in microbial alpha diversity due to drought and plant diversity loss can have a cascading effect on the plant community. The diversity of soil microbes, especially mutualists, is generally believed to enhance plant productivity through complementary ecosystem services such as litter decomposition (Hazard and Johnson (2018), Maron et al. (2018)), or by increasing the likelihood of including highly beneficial stains (Wardle (1999), Vogelsang et al. (2006)). Similarly, global change can also drive shifts in soil microbial composition that feed back to influence plant community productivity through complementarity or selection effect (Xu et al. (2025)). While dimension-reduction methods (e.g. PCA, NMDS) have been traditionally used to study microbial composition in plant diversity-productivity relationship, recent advances in bioinformatics tools (e.g. DESeq Anders et al. (2012), MaAsLin Mallick et al. (2021)) now allow for more precise identification of microbial taxa that simultaneously respond to global change and associate with plant productivity.

Here we use a biodiversity experiment in central Texas grassland to study the effect of the soil microbiome on ecosystem productivity under different scenarios of biodiversity loss and environmental contexts. We experimentally manipulated water addition, plant species richness, and plant PD, and measured plant productivity (percent cover), functional trait diversity, and trait community weighted means (CWMs). We focus on the soil fungal communities due to the ubiquitous mycorrhizal associations and their importance in plant stress responses (Tedersoo et al. (2020), Shi et al. (2023)). In our grassland system, the arbuscular mycorrhizal fungi (AMFs) are especially important for facilitating plant drought resilience and acting as a potential buffer against global changes on ecosystem functions. Thus, we paired the biodiversity experiment with soil fungal sequencing to answer the following main questions:

1. What is the effect of water availability, plant species richness, and plant PD on fungal and AMF diversity and composition?
2. What is the relationship between plant trait values and trait diversity on the diversity of different fungal guilds?
3. Do water availability and community weighted mean of plant traits mediate microbial response to plant diversity loss?
4. How do the above changes in fungal and AMF communities associate with plant productivity?

## Material and methods

### Study Site Description

The experiment was set up at the Brackenridge Field Laboratory (30°17’03.0”N 97°46’52.6”W) of University of Texas at Austin in fall 2020, with finalized species composition maintained since fall 2021 (Appendix 1.1). The study system is characterized as a central Texas grassland, subjected to constant natural drought (US drought monitor, https://droughtmonitor.unl.edu/) with mean annual temperatures of 20.9°C (January mean: 10.5°C - August mean: 30.4°C) and mean annual precipitation of 73mm (PRISM Group, Oregon State University, https://prism.oregonstate.edu, data created 4 June 2025, accessed 4 June 2025), and sandy loam river-bottom soil. The focal species of the experiment consisted of twelve native Texas cool-season grasses and wildflowers (species list see Table S1.1). Species were seeded at a constant density of 1,000 stems/m^2^ every year in November and flowered from March to June.

### Experimental Design & Covariate collection

The experiment consists of three treatments: plant species richness, plant phylogenetic diversity, and watering. In the species richness treatment, monocultures and 2-, 4-, 6-, and 12-species polycultures were planted in 2m×2m plots. The 2, 4, and 6-species richness levels were further crossed with a phylogenetic diversity (PD) treatment. The numeric values of PD (PDv) were calculated as the sum of branch lengths of all constituent species leading to the root of the phylogenetic tree (“Faith’s PD”, Faith (1992)) using the *pd* function in the ‘picante’ package (Kembel et al. (2010)). Within each of these richness level, PDv was grouped into categories (PDc) to select for plant species combinations of high, medium, and low PDc. Each diversity treatment combination received a watering (50% additional water) and an ambient rainfall (natural drought) treatment, and was replicated 3 times in 3 blocks. The three blocks were arranged to capture potential variation in soil types and management histories. In total, there were 144 plots ((3 species richness levels × 3 phylogenetic diversity levels + 12 monocultures + 3 12-species plots) × 2 watering treatments × 3 replicates). Before soil collection, we surveyed plant percent cover at each plot and measured 9 plant aboveground traits (leaf area LA, leaf mass per area LMA, leaf d15N, leaf d13C, leaf percent N, leaf percent C, leaf C:N, vegetative height, reproductive height) and 4 belowground traits (rooting depth, root tissue density RTD, specific root length SRL, root diameter). These species-level traits were sampled from the 12-species non-watered plots in 2022, and then weighed by the species relative abundance to calculate community weighted means (CWM) for all plots in both years (Ortiz and Cinoğlu et al. (in prep.)).

### Soil collection and sequencing

We collected all soil samples from one 1mx1m quadrat within each 2mx2m plot in late May 2022 and early June 2023, when plant species achieved their peak biomass. Within each quadrat, we took three subsamples from the top 5cm soil in three locations away from plants more than 5cm tall to limit the impact of any single plant species, and pooled them into one unique sample to control for fine-scale spatial variation. To minimize cross-contaminations, we sanitized the collection tools using 5% bleach, 70% ethanol, and RNAseZap between each plot. Collected samples were stored at -80° C until their DNA extraction. There were a total of 288 experimental samples, plus various negative controls and mock communities (Appendix 1.2).

We extracted DNA from the thawed soil samples using the DNeasy PowerSoil Pro Kit (Qiagen) following the manufacturer’s protocol. Library preparation and sequencing were conducted by the University of Connecticut’s sequencing center (https://core.uconn.edu/resource/mars/) following their standard procedures. Briefly, the fungal ITS2 region was amplified with fITS7 (Ihrmark et al. (2012)) and ITS4 (White et al. (1990)). Samples were amplified in triplicate 15ul reactions using Go-Taq DNA polymerase (Promega) with the addition of 3.3μg BSA (New England BioLabs), and 0.1pmol primer without the indexes or adapters to prevent inhibition from the host DNA. The ITS2 PCR reaction was incubated at 95 °C for 2 minutes, then 5 cycles of 30 s at 95.0°C, 60 s at 48.0°C and 60 s at 72.0°C, then 25 cycles of 30 s at 95.0°C, 60 s at 55.0°C and 60 s at 72.0°C followed by final extension as 72.0°C for 10 minutes. The PCR products were normalized, bead-cleaned, and sequenced on MiSeq v2 PE250 platform.

### Bioinformatics

We removed primers from demultiplexed sequences using cutadapt (Martin (2011)), merged, and clustered reads into amplicon sequencing variants (ASVs) using DADA2 (Callahan et al. (2016)). Denoising using unoise3 (Edgar (2016)) and further clustering at 97% with UCLUST performed worse at recovering the mock communities (Appendix 1.2), so we used DADA2 ASVs for all down steam analyses. We confirmed that negative controls were distinct from experimental samples in community composition (Fig. S1.3) and removed them from analyses. We rarefied reads from 8 mock community samples to 99% coverage using the coverage-based rarefaction (Chao and Jost (2012)). Rarefaction did not change the mock community recovery nor improve the weak read-abundance relationship, where sequencing reads were poorly correlated with DNA concentrations of the recovered mock community taxa (Fig. S1.2). As a result, we used non-abundance-based metrics (i.e. richness, Jaccard dissimilarity, presence-absence data) throughout the analyses.

We applied the same rarefaction procedure to the 288 experimental samples and dropped 10 samples unable to achieve 99% coverage. The remaining 278 samples were sequenced close to saturation and thus included for all subsequent analyses. We matched ASVs to known taxon using the UNITE database (Abarenkov et al. (2024)). ASVs under the phylum Glomeromycota were identified as Arbuscular Mycorrhizal Fungi (AMF).

### Statistical analysis

All analyses were conducted in R (R Core Team (2021), version 4.2.2) and visualized using ‘ggplot2’ (Wickham (2016)). To analyze the fungal response to treatments, we first tested the individual effect of watering, plant species richness, plant phylogenetic diversity value (PDv) on total fungal observed richness using generalized linear mixed-effects models with a negative binomial distribution (Warton et al. (2016), debate see: St-Pierre et al. (2018)) to account for overdispersion (*glmer.nb* in the ‘lme4’ package Bates et al. (2015)). We square-root transformed the AMF observed richness to reduce heteroscedasticity and analyzed the treatment effects using linear mixed-effects models (*lmer* in ‘lme4’). To account for spatial structure and repeated measures, all mixed effect models included “plot” nested within “block” and crossed with “year” as random effects. We then tested the effects of individual treatments on fungal and AMF community composition (Jaccard dissimilarity) using permutational analysis of variance (PERMANOVA, *adonis2* in ‘vegan’ Oksanen et al. (2025)), and followed up using *permutest* (‘vegan’) to distinguish centroid shifts from dispersion effects. Single-treatment effects were visualized via distance-based redundancy analysis (*dbrda* in ‘vegan’).

We next tested whether plant functional trait metrics predicted total fungal, AMF, and putative pathogen richness (Q2). Plant pathogen identity was inferred using FUNGuild (Nguyen et al. (2016)). Due to known limitations in amplicon-based functional inference, pathogen richness was not examined beyond its association with plant traits, which are supported by clear theoretical frameworks (Wright et al. (2004), Bergmann et al. (2020)). We modeled the scaled CWM of 13 plant traits (4 root, 9 shoot) individually as predictors of fungal and AMF richness. We also tested the effects of scaled above-and belowground functional dispersion (Laliberté and Legendre (2010a)), a widely used metric of trait diversity (Luza et al. (2023)). Trait CWMs and functional dispersion were calculated using *dbFD* in the ‘FD’ package (Laliberté et al. (2014), Laliberté and Legendre (2010b)).

To assess interactive effects between plant diversity loss and drought or plant traits (Q3), we tested the effect of watering × plant richness, watering × plant PDv, and plant PDc (categorical LMH) nested within richness on fungal and AMF richness using mixed effect models. We specifically tested if plant diversity (richness or PDv) interacted with SRL and root diameter on influencing AMF richness, or with RTD, LA, LMA, percN, percC, and C:N on influencing pathogen richness. We expect a more positive plant-soil diversity relationship under water addition and in communities with more mycorrhizal-dependent or pathogen-prone plants. We then tested the effects of water × plant richness and water × plant PDv on fungal and AMF Jaccard dissimilarity in PERMANOVA. We also assessed treatment-by-year and treatment-by-block interactions on fungal richness and composition – when interactions were significant, we ran separate mixed effect models and stratified PERMANOVAs to isolate treatment effects at each level of the interacting variable.

Lastly, we examined whether shifts in AMF richness and fungal composition predicted measures of plant productivity and biodiversity effects (Q4). Total fungal richness was excluded from this analysis, as it did not respond to treatments (Fig. 1). We first predicted plant percent cover using AMF richness and each of the experimental treatments (water, richness, or PDv). If AMF richness remains a significant predictor after accounting for treatments, this would support a non-spurious association (McElreath (2018)) between it and plant productivity. We then tested the predictive power of AMF richness on three biodiversity effect metrics – complementarity effect (CE), selection effect (SE), and total diversity effect (DE). These biodiversity effect metrics were calculated following Loreau & Hector (Loreau and Hector (2001)). Briefly, CE quantifies the average overyielding of species in polyculture relative to expected yields and measures if each monoculture species yields more when it’s in the polyculture. SE is the covariance between a species’ monoculture yield and its overyielding in polyculture and measures if the highly productive species is over-represented and drives overyielding in polycultures. DE is the sum of CE and SE. We then used MaAsLin2 Mallick et al. (2021)) to identify individual ASVs whose prevalence is significantly associated each of the three treatments (watering, plant richness, plant PDv) and five productivity metrics (plant cover, adjusted cover, CE, SE, DE). To test whether treatment-sensitive ASVs were also more predictive of productivity, we calculated Pearson correlations between their effect sizes. A significant positive correlation would suggest that fungal composition mediates plant productivity and biodiversity effects.

**Figure 1:**
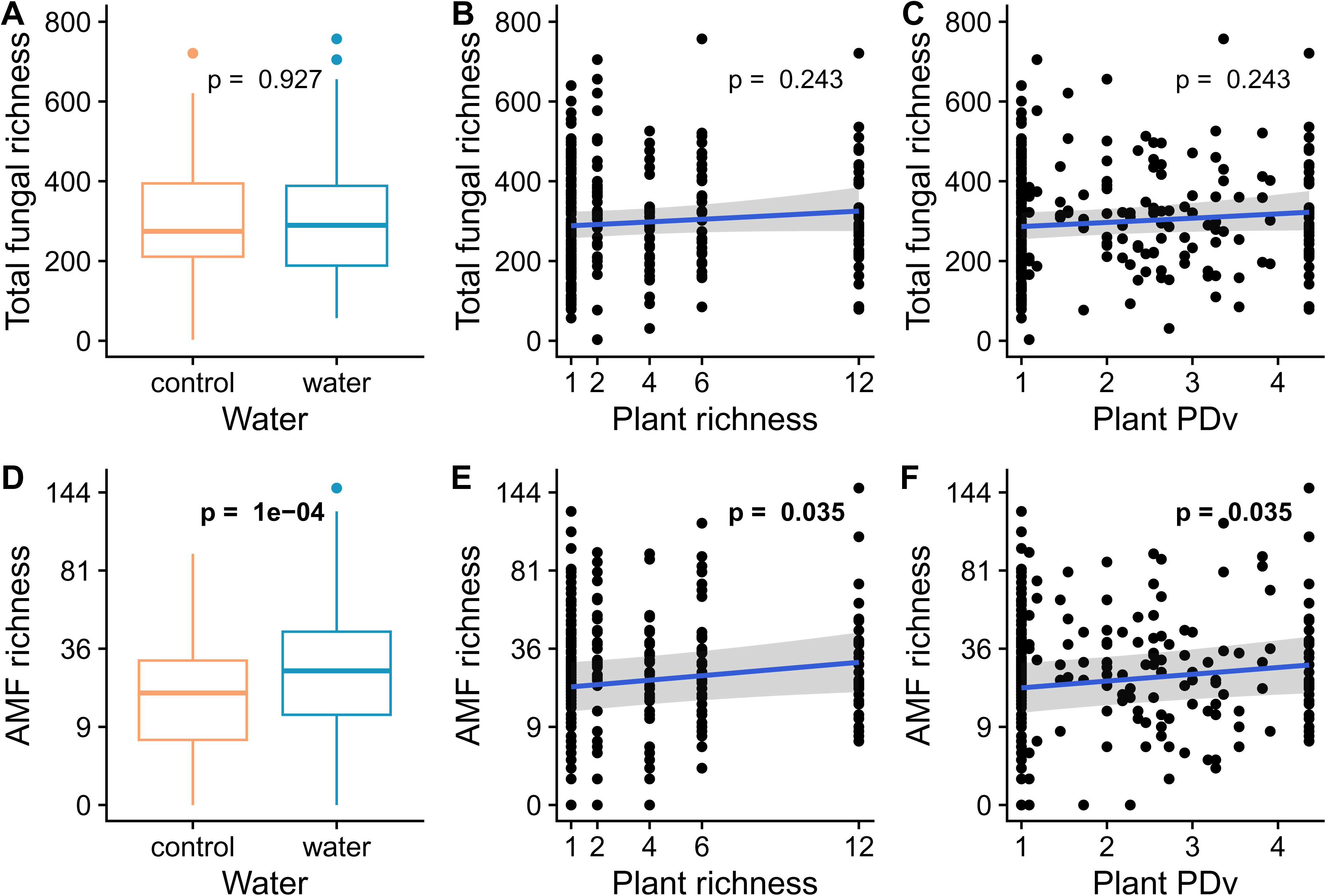
Effect of experimental treatments on fungal richness (A-C) and square-root-transformed AMF richness (D-E). Fungal richness did not differ in watered versus ambient plots (A), did not increase with either targeted plant richness (B) or targeted plant phylogenetic diversity (C). AMF richness was higher in watered plots (D), and plots with higher targeted plant richness (E) or PDv (F). There were no significant interactive effects between water addition and plant diversity (either richness or PDv). Predicted trend lines, 95% CIs, and p-values were based on the underlying (generalized) linear mixed effect models.

In all analysis above (except for excluding spurious correlations), we tested each explanatory variable individually to understand its specific influence. We presented raw p-values of each individual model in the main text, and adjusted their p-values using the Benjamini–Hochberg (Benjamini and Hochberg (1995)) procedure in the Appendix.

## Results

We identified 14278 amplicon sequencing variants (ASVs) contained in 278 experimental samples, including 2814 singletons. There were 1928 AMF ASVs, including 251 singletons.

### 1. Independent effect of plant diversity and water addition on AMF richness

Watering, plant richness, and plant PDv treatment did not significantly alter the total fungal richness (Fig. 1A-C; all p > 0.1). On the other hand, all three treatments increased the total AMF richness (Fig. 1D-E; water: β = 1.19, p < 0.0001; plant richness: β = 0.086, p = 0.035; plant PDv: β = 0.26, p = 0.031). The low, medium, and high PDc treatment nested within each richness was not a significant predictor of either fungal or AMF richness (both p > 0.1). There was no significant interactive effect between water and either one of the plant diversity measures on either the total fungal richness or the AMF richness (Table S2.1; all p > 0.1). However, in explaining AMF richness, the model that contained both water addition and plant diversity had R² (marginal R²: water + richness = 0.079; water + PD = 0.080) close to the sum of R² from models with individual treatment effects (marginal R²: water = 0.062; richness = 0.018: PD = 0.018). There was no significant interactive effect between year and any treatments on either total fungal or AMF richness (Table S2.2), meaning the treatment effect on fungal richness was consistent throughout both years.

### 2. Orthogonal effect of plant diversity and water addition on soil fungal composition

Water, plant richness, and plant PDv influenced the total fungal composition (all p < 0.001), but these differences in fungal communities were all driven by different dispersions (all p <0.001). Plant PDc within richness again had no significant effect on total fungal compositions (p > 0.1). Similarly, plant richness and PDv, but not PDc within richness, significantly influenced AMF composition (p = 0.016/0.035/0.76) and dispersions (p < 0.001 for both plant richness and PDv). Watering also had a significant effect on AMF composition (p < 0.001), and the separation was driven by different centroids than dispersions (p = 0.072). However, despite the significant effect of the treatments, each of them only explained less than 1% the total variance explained in fungal composition. Similarly, in the complementary analysis using distance-based redundancy analysis (dbRDA), the three treatments combined accounted for less than 2% of the total variance in both total fungal and AMF communities. There was no significant interactive effect between water addition and either plant diversity measure (plant richness or PD) on either total fungal or AMF richness (all p > 0.1). Likewise, in the dbRDA plots for total fungal and AMF communities (fig 2), the vectors corresponding to the watering effect were orthogonal to those corresponding to the plant diversity (richness and PD) effect, indicating relatively independent effects of water and plant diversity on soil fungal compositions. All three treatments significantly interacted with year and block, but redoing the PERMANOVA by permuting within strata of year or block did not qualitatively change our results above.

**Figure 2:**
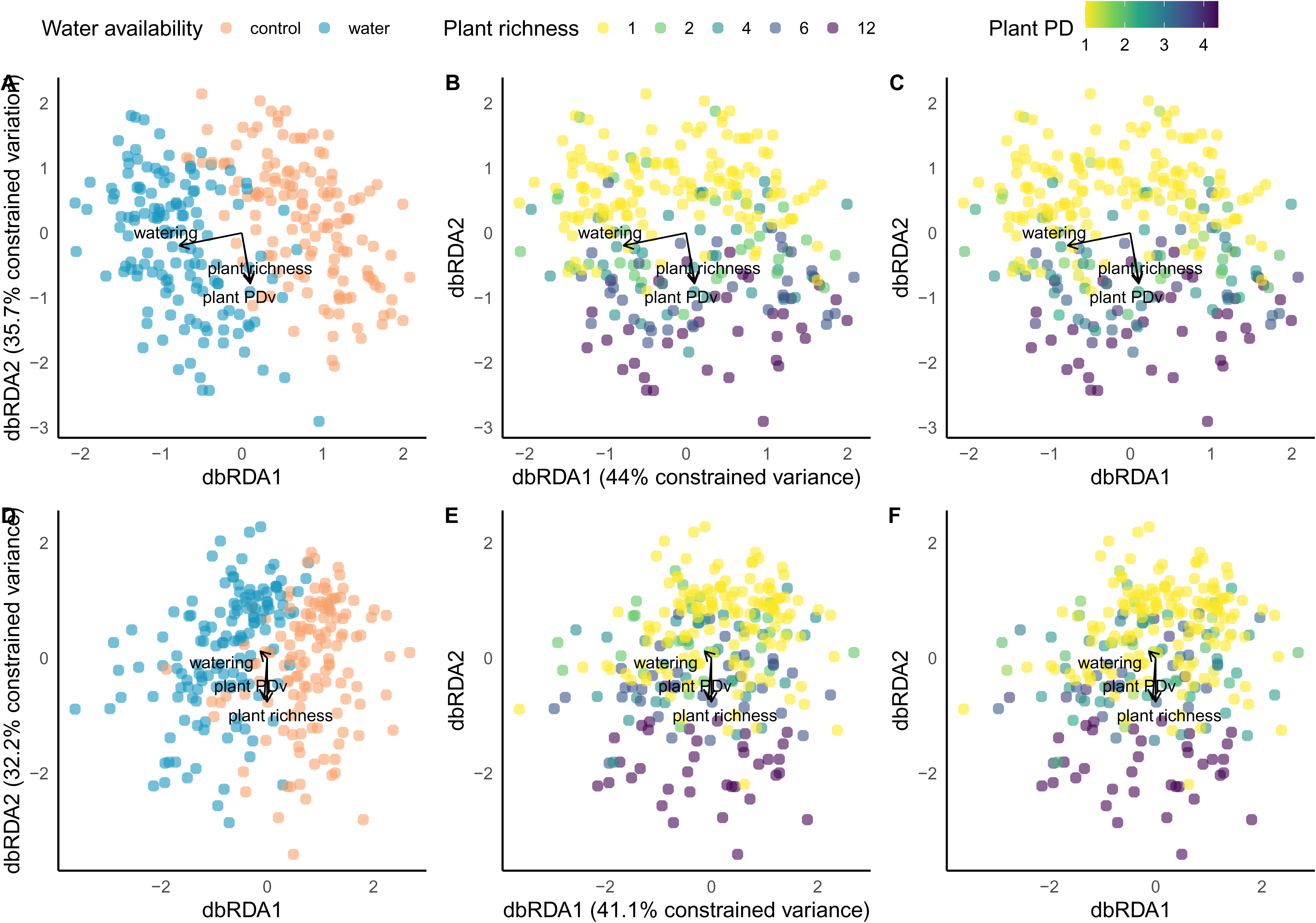
Effect of experimental treatments on fungal (A-C) and AMF (D-F) composition with Jaccard-distance-based redundancy analysis (dbRDA). Fungal community dissimilarity were significantly separated by water (A; PERMANOVA p<0.001) along the first RDA axis, and separated by plant richness (B; PERMANOVA p<0.001) and PDv (C; PERMANOVA p<0.001) along the second RDA axis. The constrained variance (explained by three treatments) in fungal composition is 1.7% of the total variance in fungal composition. AMF community dissimilarity were separated by water (D; PERMANOVA p<0.001) mostly along the first RDA axis, and separated by plant richness (E; PERMANOVA p=0.016) and PDv (F; PERMANOVA p=0.035) along the second RDA axis. The constrained variance in fungal composition is 1.3% of the total variance in fungal composition.

### 3. Trait diversity and different CWMs of plant traits explain AMF and pathogen alpha diversity

Richness of various fungal groups generally positively correlates with functional trait diversity. Total fungal richness was positively correlated with below-ground functional dispersion but not significantly correlated with above-ground functional dispersion (Fig. 3; Table S2.3). Both above- and below-ground functional dispersion positively correlates with AMF richness and potential soil pathogen richness (Fig.3; Table S2.3).

**Figure 3:**
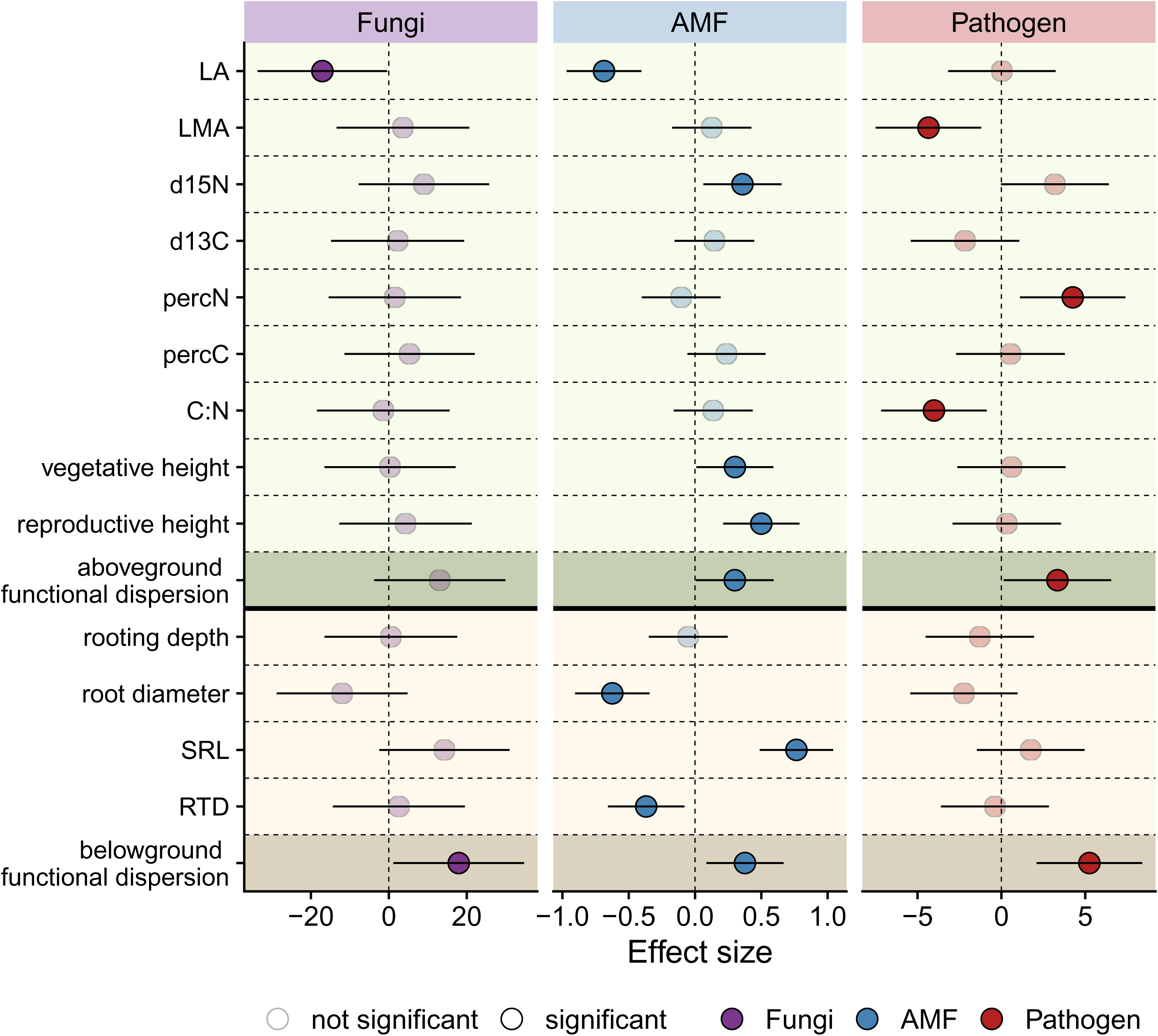
Effect of scaled plant above-ground and belowground trait community weighted means (CWMs) and functional dispersions, on total fungi, AMF, and plant pathogen richness in soil. Total fungi and pathogen richness were not transformed, and AMF richness was square-root-transformed. Circles to the right of zero (dashed line) indicate positive correlations and those to the left of zero indicate negative correlations. Dark circles have 95% CI not overlapping zero and light circles have 95% CI overlapping zero. Aboveground and belowground trait CMWs were respectively shown with light green and light tan backgrounds. Aboveground and belowground trait functional diversity were respectively shown with darker green and darker tan background.

No individual plant trait CWMs significantly correlated with the total fungal richness, except CWM of leaf area (Appendix), but different traits can help predict AMF or pathogen richness. For belowground traits, AMF richness was positively associated with specific root length (CWM, same for other traits; β = 0.76, p < 0.0001) and negatively associated with root diameter (β = -0.62, p < 0.0001) and root tissue density (β = -0.37, p = 0.012). For aboveground traits, AMF richness negatively correlated with leaf area (β = -0.69, p < 0.001) and positively correlated with leaf d15N (β = 0.36, p = 0.017). Moreover, AMF richness was also positively correlated with both plant vegetative (β = 0.30, p = 0.043) and reproductive heights (β = 0.50, p = 0.001). On the other hand, the richness of potential soil pathogen was positively associated with leaf percent N (β = 4.25, p = 0.008), and negatively associated with both LMA (β = - 4.35, p = 0.007) and leaf C:N (β = - 4.02, p = 0.012), all indicating higher pathogen richness in “faster” communities. However, these traits were no longer significant predictors of pathogen richness when we only included probable and highly probable plant pathogens (Fig. S2.1), or after BH adjustment (Fig. S2.2, Table S2.3). Contrary to expected, plant richness and PDv did not significantly interact with the CWM of either SRL or root diameter in influencing the AMF richness; similarly, the effect of plant diversity on soil pathogen richness did not depend on the CMWs of any “fast-slow” plant traits: RTD, LA, LMA, percN, percC, or C:N (Table S2.4).

### 4. Soil fungal communities are correlated with plant productivity and diversity effects

AMF richness was positively correlated with plant percent cover, a proxy for productivity. This relationship remained significant even after accounting for the water addition treatment (Fig. 4A), plant richness (Fig. 4B), or plant PD (Fig. 4C) as confounding variables. These findings reduce the likelihood that the observed relationship between AMF richness and plant productivity was a spurious outcome of shared responses to treatment. AMF richness was also positively associated with selection effect in the plant community (Fig. 4E; p = 0.022), but not significantly linked to complementarity (Fig.4D; p = 0.85; significant negative association after removal of an outlier: p = 0.004) or net biodiversity effect (Fig. 4F).

**Figure 4:**
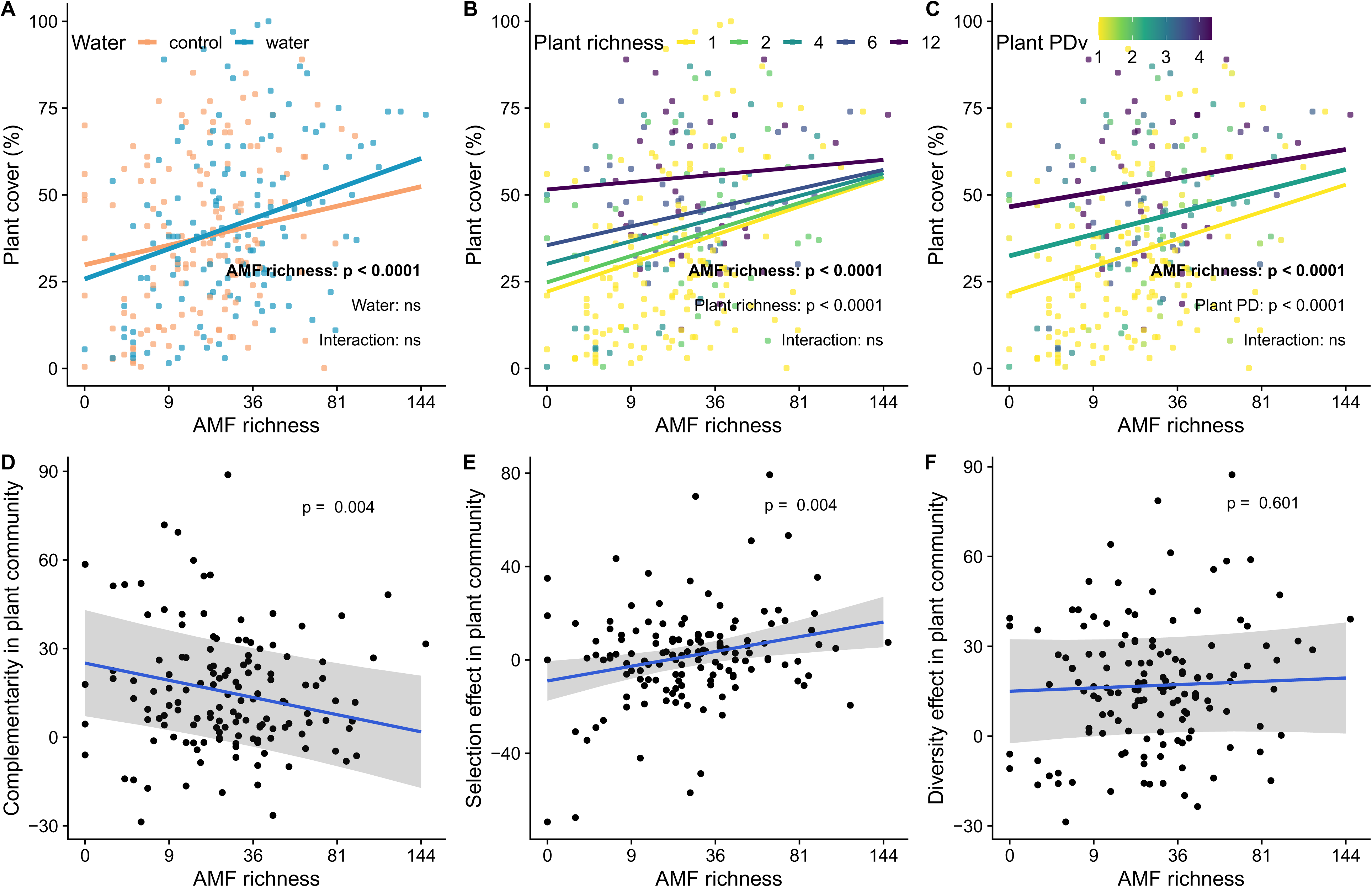
Correlation between AMF richness (all square-root-transformed) and plant productivity (A-C) and biodiversity effects (D-F). AMF richness was significantly and positively correlated with plant percent cover even after accounting for water aaddition (A), plant richness (B), or plant PD (C), demonstrated by the positive slope for all levels. AMF richness was negatively correlated with complementarity in plant communities when an outlier was removed (as shown in D), but the correlation was no longer significant with the outlier present (Fig. S2.3). AMF richness was positively correlated with selection effect (E) and not correlated with net diversity effect (F) regardless of the presence of the outlier.

Similar to AMF richness, fungal composition also plays a potential role in mediating plant biodiversity-ecosystem-functioning relationships. The effect size of water addition on fungal ASV prevalences was not significantly correlated with the effect size of plant cover on fungal prevalences, whether or not water addition was included as a confounding covariate (Fig. 5 correlation plot top left region; water ∼ cover: r = 0.005, p = 0.89; water ∼ cover|water: r = -0.06, p = 0.099). In contrast, both plant richness and PDv showed strong positive correlations between their effects on ASVs and the effect of plant cover on ASVs (Fig. 5 correlation plot top right region; plant richness ∼ cover: r = 0.63; plant richness ∼ cover|richness: r = 0.44; plant PDv ∼ cover: r = 0.66; plant PDv ∼ cover|PDv: r = 0.46; all p < 0.0001). Furthermore, watering and plant diversity treatments also showed divergent effects on driving plant diversity effects through fungal ASVs (Fig. 5 correlation plot bottom region). The extent to which ASVs were driven by water addition was positively correlated with the extent to which they were associated with the selection effect (r = 0.13, p = 0.001) and negatively correlated with their associations with both the complementarity (r = -0.24, p < 0.0001) and the net diversity effect (r = -0.19, p < 0.0001). Conversely, the extent to which ASVs were driven by plant richness or PDv was not significantly correlated with the extent to which they were associated with the selection effect (plant richness ∼ selection: r = -0.01, p = 0.74; plant PDv ∼ selection: r = -0.04, p = 0.25) but positively correlated with their associations with both the complementarity (r = -0.24, p < 0.0001) and the net diversity effect (r = -0.19, p < 0.0001).

**figure 5:**
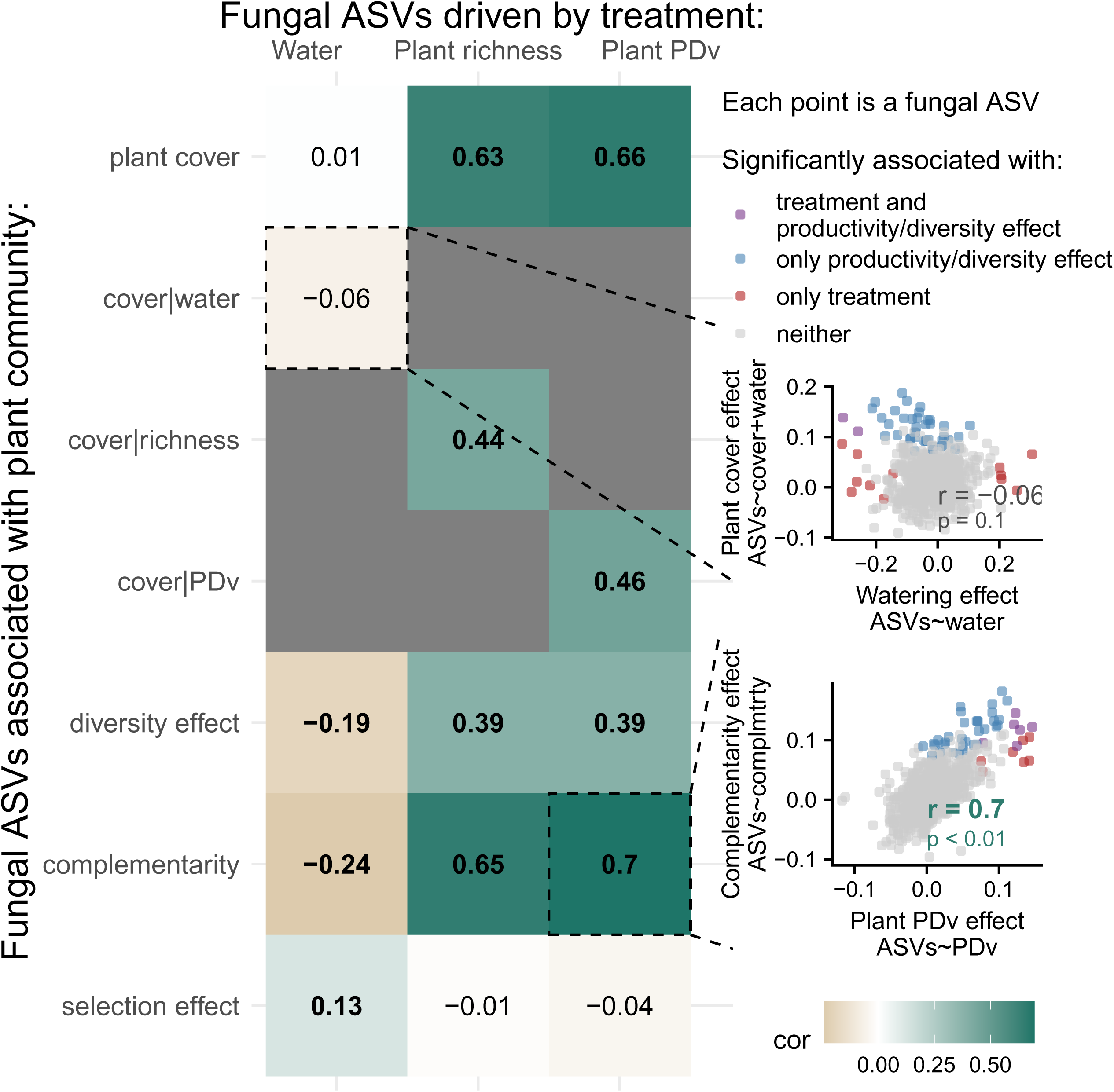
Water and plant diversity treatments exhibited divergent effects on plant community productivity and diversity effect via fungal ASVs. Left Panel shows the Pearson correlation between total fungal ASVs driven by treatments and those associated with various plant community measure. Each value measures the correlation between the effect sizes of two multivariate associations – one between ASVs and one of the three treatments, and one between ASVs and one of the six productivity measures. Bolded values indicate significant (p < 0.05) correlations. For example, the Top Right Panel shows a non-significant correlation (r = -0.06) between the effect sizes for ‘water’ in the regression ‘ASVs ∼ water’, and the effect sizes for ‘plant cover’ in the regression ‘ASVs ∼ plant cover + water’ after accounting for the potential confounding effect of water. In other words, each point represents an ASV; its location on the x axis indicates how much this ASV associated with water addition, and its y axis location indicates how much it associated with plant cover after accounting for water addition. Similarly, the Bottom Right Panel is another example of a significant positive correlation (r = 0.7) between how much the ASVs were driven by the plant PD treatment, and how much they were associated with complementarity in the plant community.

## Discussion

Our study provides an integrated assessment of soil microbial communities under global change and their role in plant community feedback and biodiversity-ecosystem functioning relationships. First, water addition and plant diversity (species richness or PDv) independently increased soil AMF richness and altered soil fungal composition. Moreover, the functional composition of soil fungal communities and their richness can also be explained by plant functional trait CWMs and functional trait diversity. However, the plant-soil diversity relationships did not depend on either water availability or plant trait CWMs. Lastly, we applied an underutilized method in this field to show that AMF richness and total fungal composition, now both threatened by drought and plant diversity loss, were indicative of plant productivity potentially through selection effect or complementarity. Together, these results suggest that plant diversity loss and drought impact the soil fungal communities with cascading effects on ecosystem productivity, as these soil microbiomes can mediate plant diversity-productivity relationships via multiple pathways in our grassland ecosystem.

Water addition, plant richness, and plant PDv all increased AMF richness but did not change total soil fungal richness (Fig. 1). The treatment effects on AMF richness were consistent with previous findings that AMF richness tends to decrease with drought (Lozano et al. (2021)) and increase with plant diversity (Fahey et al. (2023)). The differential response of AMF vs total fungal communities to plant diversity can be partially explained by AMF, an obligate mutualist, having a tighter interaction with the plant communities than the total fungal community in the bulk soil, which contains free-living fungi such as saprotrophs (Millard and Singh (2010)). This was in line with the global pattern that plant diversity does not explain grassland fungal alpha diversity (Prober et al. (2015)). Similarly, the sensitivity of AMFs compared to other fungal taxa to water addition may also be explained by their symbiosis with plants (Fu et al. (2022)). However, it was also possible that the sequencing noise and error were higher in the total fungal community than the characterized AMF communities. The positive relationship between plants functional diversity and richness of total fungi and AMF (Fig. 3) was consistent with the positive effect of plant species richness and phylogenetic diversity and supported the general “diversity-begets-diversity” hypothesis (Madi et al. (2020)). While functional diversity was not manipulated as a treatment, but rather a covariate or even outcome of our plant diversity treatments, our results nevertheless showed that loss in plant diversity not only directly affected fungal richness and composition, but also could indirectly do so through decline in plant functional diversity.

The lack of interactive effects between plant diversity (richness or PDv) and water addition on both fungal and AMF communities suggested the effect of biodiversity loss and drought on fungal communities might not depend on one another in our system (versus interactive effect in other systems, Xi et al. (2022)). In fact, the effect of water and plant diversity on AMF richness seem to be additive (additive 𝑅^2^), and their influence on total fungal and AMF composition was independent (Fig. 2), both indicating that abiotic and biotic factors may affect fungal community through separate mechanisms – e.g. drought can select for stress tolerant fungi (Malik et al. (2020), Treseder and Lennon (2015)), whereas plant diversity and identity may select for host association (Plett and Martin (2018)) and resource acquisition traits (Malik et al. (2020), Treseder and Lennon (2015)) in fungi. Thus, drought impacts on fungal communities are unlikely to be mitigated by higher plant diversity, and conversely, the effects of plant diversity loss may not be offset by water addition either. Within plant diversity, plant species richness and phylogenetic diversity had highly similar effects on the soil microbial communities. However, although AMF and fungal communities were influenced by PD across all richness levels (PDv), they did not differ across the low, medium, and high PD levels (PDc) within each richness level. This implies that the effect of plant diversity on fungal communities may only be evident at broader evolutionary scales, as closely related plant species may share the same generalist fungal symbionts (Öpik et al. (2009), but see Reinhart and Anacker (2014)).

Additionally, none of these treatment effects on total fungal/AMF richness depended on year as contrary to expected, despite the total fungal and AMF richness themselves and many environmental covariates (rainfall, plant productivity) differing across the two years. This may indicate the relative robustness of soil fungal communities (Gschwend et al. (2021)) and indirectly corroborate the strong individual, but not interactive, treatment effects on the soil fungal communities.

How and why plant traits may affect the richness of different fungal guilds can be understood through the lens of the leaf economic spectrum (Wright et al. (2004)) and root economic space (Bergmann et al. (2020)). Usually, the “collaboration gradient” in the root economic space offers insights on patterns of plant-mycorrhizal associations, whereas the fast-slow “conservation gradient” is used to understand plant-pathogen interactions. Surprisingly, we found that AMF richness was positively correlated with less “outsourcing” strategies of the plant community (indicating resource acquisition by roots and less mycorrhizal dependent; Fig. 3), although we would otherwise expect more mycorrhizal dependent plants to foster more, and more diverse, AMF mutualists. However, previous studies have shown similar positive relationships (Wu et al. (2025)), no relationships (Hennecke et al. (2023)), or negative relationships (Hennecke et al. (2025)) between AMF richness or diversity and less outsourcing plant strategies. The various possible relationships imply either that AMF richness is not a direct indicator of mycorrhizal dependence of the plant (but see Ferreira et al. (2021)), or that plant-AMF association is complex and context dependent (Romero et al. (2023)). On the other hand, higher richness of potential soil pathogens generally co-occurred with a different set of, but unanimously “faster”, traits (lower LMA, lower leaf C:N; Fig. 3). This is in line with many previous studies (Cappelli et al. (2020), Coley et al. (1985)) that suggest higher natural enemy occurrence in “fast” plant communities due to weaker defenses. However, pathogen-trait correlations tend to be less robust than AMF-trait correlations, likely due to the context dependence of soil pathogens and the ongoing challenge of characterizing pathogens from amplicon sequencing.

The relationship between fungal functional guilds and CWMs of plant traits altogether revealed four new insights. First, the different responses of fungal guilds to the CWM of different plant traits could partially explain why total fungal richness did not correlate with the CWM of most traits, and suggest that plant functional traits capture distinct plant physiological processes that shape different fungal guilds. Therefore, shifts in plant community strategy due to global change (Kimball et al. (2016), Griffin-Nolan et al. (2019)), even without biodiversity loss, could change the soil fungal communities and their downstream effects. While amplicon sequencing and current databases remain imperfect for functional assignment of fungal pathogens (Tedersoo et al. (2022)), our results regardless provided valuable insights for hypothesis generation. Future trans-omics studies can identify candidate genes, pathways, and microbial traits (Malik et al. (2020)) that underpin these divergent microbial responses to plant traits.

Second, contrary to expected, the diversity and CWM of above-ground traits can explain soil AMF and pathogen richness as well as those of below-ground traits. While root traits have traditionally been regarded as the main mirror of plant-soil-microbe interactions, these interactions are a holistic process that involves various aspects of plant physiology. For example, plant above ground traits can explain soil microbial composition and function through plant biomass, litter quality, and soil abiotic conditions (Legay et al. (2014), Grigulis et al. (2013)); on the other hand, soil microbes can in turn affect plant growth, phenology, and above ground traits (Lau and Lennon (2011)). Third, both functional diversity and CWM of traits affected AMF and pathogen richness, hinting at both “complementarity” and “selection effect” in fungal community assembly – AMF and pathogen richness can increase with plant functional diversity via niche partitioning and reduced competition, or with certain plant traits that disproportionately foster diverse fungal communities. Lastly, the plant-soil diversity relationship in our system did not depend on dominant plant strategies, similar to the way it did not depend on water availability. Therefore, the negative effect of plant diversity loss might consistently impact different plant communities and could not be mediated through plant community turnover or trait plasticity.

AMF and soil microbes not only respond to changes in the plant communities, but can also affect processes aboveground such as mediating the plant diversity-productivity relationship (Klironomos et al. (2000), Hazard and Johnson (2018)). We found AMF richness positively correlated with plant cover, even after accounting for the fact that they were both driven by the treatments (Fig. 4 A-C). The robust positive relationship after controlling for the confounding variable largely precluded the possibility of spurious correlation. Therefore, the decline in AMF richness due to drought or plant diversity loss may drive a decrease in plant productivity (Van Der Heijden et al. (1998), Vogelsang et al. (2006)); or alternatively, decrease in plant productivity due to plant diversity loss may drive a decline in AMF richness (Cline et al. (2018)). However, AMF richness was positively associated with selection effect but not significantly associated with complementarity (Fig. 4D-E), contrary to the popular hypothesis that diverse mutualists increase complementarity by reducing niche overlaps in plant communities (Hazard and Johnson (2018), but see Wagg et al. (2015)). Although counterintuitive, similar patterns have been often observed (Wagg et al. (2011b), Wagg et al. (2011a), Hazard and Johnson (2018)) and could be attributed to a more effective sampling and inclusion of more beneficial AMF strains (Wardle (1999), Vogelsang et al. (2006)). In addition to AMF richness, total fungal composition could be another mediator between plant diversity and productivity. The fungal ASVs driven by plant diversity treatments were also positively correlated with cover, residual cover, complementarity and net diversity effect in the plant community (Fig. 5, columns under Plant richness and PD), indicating the effect of biodiversity loss may be connected to decrease in plant productivity through shifts in fungal composition. On the other hand, watering had no or opposite effect (Fig. 5, column under Water), again showing the divergent effect of water versus plant diversity treatment on downstream community processes through the fungal community. However, these interpretations should be considered in light of the limitations of observational data. Future experiments that manipulate AMF richness or fungal composition will provide definitive insights into their effects on plant productivity and other ecosystem functioning.

## Conclusion

Potential role of soil fungi in plant diversity-productivity relationships Drought and plant diversity loss are common global change scenarios and both can lower soil AMF richness and shift the soil fungal composition. The decline in AMF richness and the shifts in individual fungal ASVs due to global change may further go hand in hand with a decline in plant community productivity, an important ecosystem function. These changes in belowground communities and their subsequent consequences could be similarly likely when plant species are lost randomly or in phylogenetically and functionally related clusters. Furthermore, the effects of water availability and plant diversity were independent of each other, highlighting the importance of simultaneously managing both drought and biodiversity loss to prevent cascading effects on ecosystems through the soil microbiome.

