## Appendix for "Plant diversity and water regime have distinct effects on soil fungal communities that are linked to plant productivity"

### Appendix 1: Supplementary Materials, Methods, and Confirmatory Analyses

#### 1.1 Experimental Design and Species list

The experiment was initially set up in fall 2020, when soil was tilled. Seeds were first sown in February 2021 (to avoid a winter storm) along with the watering treatment. However, weeding efforts that year were insufficient to maintain the intended plant richness and PD treatments, and one of the twelve originally chosen species (Prickly poppy: *Argemone albiflora*) was replaced in 2022 by *Phacelia congesta*. Beginning in fall 2021, experimental treatments were finalized – In November, all plots were reseeded and watering treatments were applied, and weeding successfully maintained the target plant diversity levels. Data analyzed in this study were collected during the 2022 and 2023 growing seasons.

In addition to the plant richness, plant PD, and water addition treatments, we used a split-plot design to test the effect of insect herbivores and pollinators. That is out of the scope of the current analysis, but briefly, we have 432 total treatments of those 144 plots split into open quadrats, insect exclusion quadrats, and shading control quadrats.

**Figure S1.1**: Schematics of the experiment.

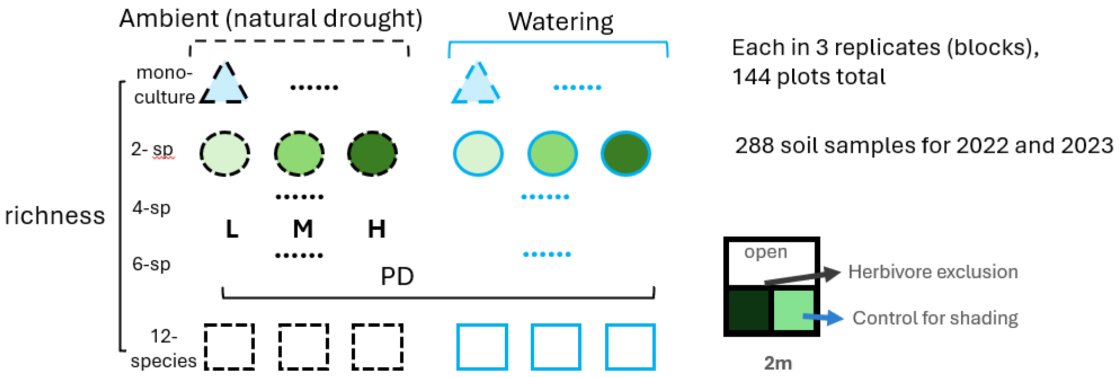

**Table S1.1**: Table of the twelve experimental plant species.

| Species | Family | Common Name | Duration | Growth form |
| --- | --- | --- | --- | --- |
| *Aristida purpurea* | Poaceae | Purple three-awn | Annual/Perennial | Grass |
| *Bouteloua curtipendula* | Poaceae | Sideoats grama | Perennial | Grass |
| *Bouteloua gracilis* | Poaceae | Blue grama | Perennial | Grass |
| *Lespedeza texensis* | Fabaceae | Bluebonnet | Annual | Legume |
| *Desmanthus illinoensis* | Fabaceae | Illinois bundleflower | Perennial | Legume |
| *Monarda punctata* | Lamiaceae | Spotted beebalm | Annual/Biennial/ Perennial | Forb/herb |
| *Monarda citriodora* | Lamiaceae | Horsemint | Annual/Biennial/ Perennial | Forb/herb |
| *Gaillardia pulchella* | Asteraceae | Indian blanket | Annual/Biennial/ Perennial | Forb/herb |
| *Coreopsis tinctoria* | Asteraceae | Golden tickseed | Annual/Biennial/ Perennial | Forb/herb |
| *Chrysopsis americana* | Asteraceae | American basket-flower | Annual | Forb/herb |
| *Ipomopsis rubra* | Polemoniaceae | Standing cypress | Biennial | Forb/herb |
| *Phacelia congesta* | Hydrophyllaceae | Blue curls | Annual/Biennial | Forb/herb |

**1.2 Negative controls, mock communities, and relevant analyses**

In addition to the 288 experimental samples, we also included 10 negative controls and 8 mock communities in sequencing across 4 plates. There were 4 extraction negatives (extracting from field blanks), 2 PCR negatives (PCR-grade water), and 4 mock community negatives (extraction negatives during the mock community preparation). There were 4 even and 4 tiered mock communities, which had equal or tiered concentration of the 32 fungal taxa (Table S1.2). After sequencing and bioinformatics, sequences appeared in mock community samples (Query Sequence) were compared with known sequences of the 32 taxa (Subject Sequence) using Nucleotides BLAST (https://blast.ncbi.nlm.nih.gov/).

To test if the number of reads of fungal sequences can be used to approximate fungal abundance, linear regressions test if the number of reads of the matched ASVs in the tired mock community are good predictors of their recorded DNA concentration. This procedure was repeated before and after rarefaction (Figure S1.2 AC versus BD), and with or without an outlier(Figure S1.2 AB versus CD).

Fungal composition of all samples was visualized on an NMDS plot, to confirm that the mock community and negative controls were distinct from the experimental communities (Figure S1.3)

**Table S1.2**: Information about the 32 fungal taxa contained in the mock community.

| Phylum | Subphylum | Class | Subclass | Order | Family | Genus | Species |
|---|---|---|---|---|---|---|---|
| Ascomycota | Pezizomycotina | Sordariomycetes | Hypocreomycetidae | Hypocreales | | Acremonium | *Acremonium kiliense* |
| Basidiomycota | Agaricomycotina | Agaricomycetes | Agaricomycetidae | Agaricales | Psathyrellaceae | Panaeolina | *Panaeolina foenisecii* |
| Chytridiomycota | | Chytridiomycetes | | Chytridiales | Chytridiaceae | Chytridium | *Chytridium confervae* |
| Ascomycota | Pezizomycotina | Leotiomycetes | Leotiomycetidae | Helotiales | Sclerotiniaceae | Botryotinia | *Botryotinia sp.* |
| Ascomycota | Pezizomycotina | Sordariomycetes | Xylariomycetidae | Amphisphaeriales | Discosiaceae | Seimatosporium | *Seimatosporium lonicerae* |
| Ascomycota | Pezizomycotina | Sordariomycetes | Sordariomycetidae | Sordariales | Coniochaetaceae | Coniochaeta | *Coniochaeta prunicola* |
| Basidiomycota | Agaricomycotina | Agaricomycetes | | Polyporales | Sparassidaceae | Sparassis | *Sparassis crispa* |
| Ascomycota | Pezizomycotina | Sordariomycetes | Hypocreomycetidae | Hypocreales | Hypocreaceae | Hypocrea | *Hypocrea orientalis* |
| Ascomycota | Pezizomycotina | Dothideomycetes | Dothideomycetidae | Capnodiales | Cladosporiaceae | Cladosporium | *Cladosporium sphaerospermum* |
| Basidiomycota | Ustilaginomycotina | Exobasidiomycetes | | Malasseziales | | Malassezia | *Malassezia* |
| Ascomycota | Pezizomycotina | Sordariomycetes | Xylariomycetidae | Amphisphaeriales | Pestalotiopsidaceae | Pestalotiopsis | *Pestalotiopsis chamaeropis* |
| Ascomycota | Pezizomycotina | Dothideomycetes | Pleosporomycetidae | Pleosporales | Pleosporaceae | Epicoccum | *Epicoccum nigrum* |
| Ascomycota | Pezizomycotina | Eurotiomycetes | Chaetothyriomycetidae | Phaeomoniellales | | Neophaeomoniella | *Neophaeomoniella zymoides* |
| Basidiomycota | Agaricomycotina | Agaricomycetes | Agaricomycetidae | Boletales | Suillaceae | Suillus | *Suillus brevipes* |
| Ascomycota | | Ascomycetes | | | | Lecanicillium | *Lecanicillium muscarium* |
| Ascomycota | Pezizomycotina | Sordariomycetes | Hypocreomycetidae | Glomerellales | Glomerellaceae | Colletotrichum | *Colletotrichum tropicale* |
| Ascomycota | Pezizomycotina | Sordariomycetes | Hypocreomycetidae | Hypocreales | Hypocreaceae | Trichoderma | *Trichoderma theobromicola* |
| Zygomycota | Mucoromycotina | | | Mucorales | Mucoraceae | Mucor | *Mucor endophyticus* |
| Ascomycota | Pezizomycotina | Sordariomycetes | Xylariomycetidae | Xylariales | Xylariaceae | Xylaria | *Xylaria mali* |
| Ascomycota | Pezizomycotina | Dothideomycetes | Pleosporomycetidae | Pleosporales | | Phoma | *Phoma sp. A cf. aliena* |
| Ascomycota | Pezizomycotina | Dothideomycetes | Pleosporomycetidae | Pleosporales | Didymosphaeriaceae | Dendrothyrium | *Dendrothyrium variisporum* |
| Ascomycota | Taphrinomycotina | Schizosaccharomycetes | Schizosaccharomycetidae | Schizosaccharomycetales | Schizosaccharomycetaceae | Schizosaccharomyces | *Schizosaccharomyces pombe* |
| Basidiomycota | Ustilaginomycotina | Ustilaginomycetes | | Ustilaginales | Ustilaginaceae | Ustilago | *Ustilago cynodontis* |
| Ascomycota | Pezizomycotina | Sordariomycetes | Hypocreomycetidae | Hypocreales | Nectriaceae | Fusarium | *Fusarium torulosum* |
| Zygomycota | Mucoromycotina | | | Mortierellales | Mortierellaceae | Mortierella | *Mortierella alpina* |
| Basidiomycota | Agaricomycotina | Agaricomycetes | | Russulales | Hericiaceae | Hericium | *Hericium abietis* |
| Zygomycota | Entomophthoromycotina | | | Entomophthorales | Ancylistaceae | | *Conidiobolus* |
| Zygomycota | Kickxellomycotina | | | Dimargaritales | Dimargaritaceae | Dimargaris | *Dimargaris bacillispora* |
| Basidiomycota | Pucciniomycotina | | | | | | *Pucciniomycotina* |
| Basidiomycota | Agaricomycotina | Tremellomycetes | Tremellomycetidae | Tremellales | Tremellaceae | Tremella | *Tremella aurantia* |
| Ascomycota | Saccharomycotina | Saccharomycetes | Saccharomycetidae | Saccharomycetales | Metschnikowiaceae | Metschnikowia | *Metschnikowia agaves* |

**Figure S1.2**: Read-abundance relationship of the tiered mock communities, using raw (A, B) and rarefied (C, D) sequence reads. In both cases, the linear relationships were mostly driven by a high-read outlier (A, C). When this outlier was removed, the relationship was no longer significant (B, D).

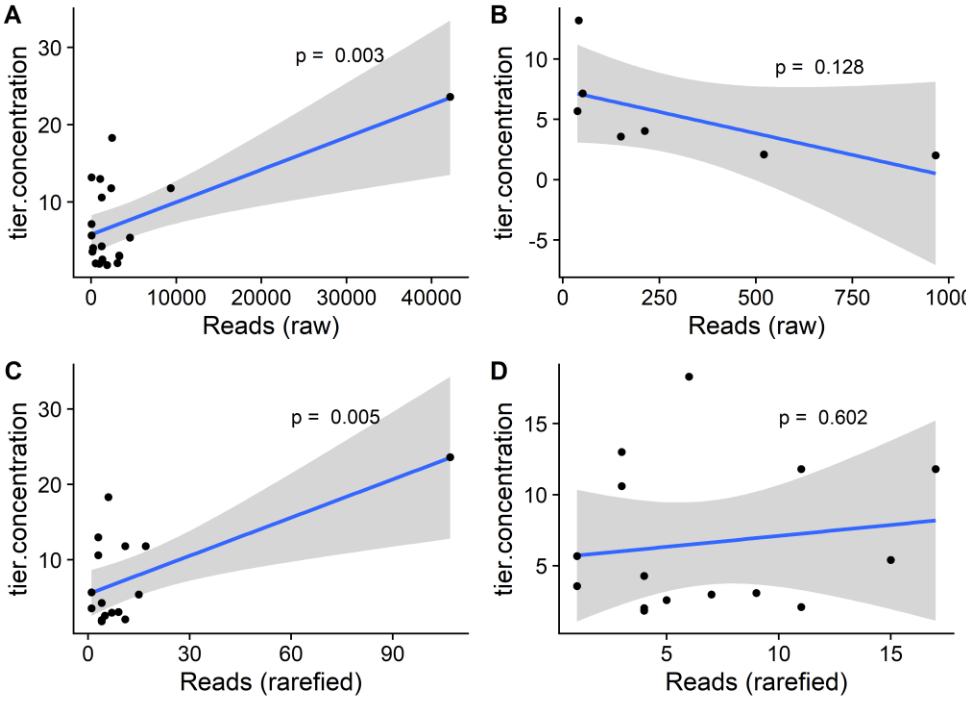

**Figure S1.3**: NMDS plot for fungal composition across different samples. Fungal composition from all experimental samples were similar to each other, whereas the negatives and mock communities form distinct clusters.

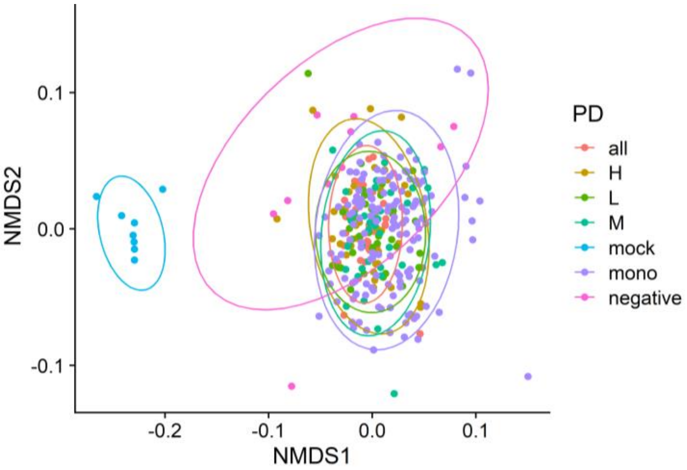

### Appendix 2: Supplementary Results

**Table S2.1**: Interaction between water regime and plant diversity treatments on total fungal and AMF richness. None of the interactions were statistically significant.

| Response variable | Interaction tested | $\chi^2$ | df | p.val |
|---|---|---|---|---|
| Fungal richness | plant richness × water | 0.54 | 1 | 0.46 |
| | plant PDv × water | 0.93 | 1 | 0.33 |
| AMF richness | plant richness × water | 1.17 | 1 | 0.28 |
| | plant PDv × water | 1.50 | 1 | 0.22 |

**Table S2.2**: Interaction between treatments and year on total fungal and AMF richness. None of the interactions were significant.

| Response variable | Interaction tested | $\chi^2$ | df | p.val |
|---|---|---|---|---|
| Fungal richness | water × year | 0.07 | 1 | 0.79 |
| | plant richness × year | 0.54 | 1 | 0.46 |
| | plant PDv × year | 0.20 | 1 | 0.66 |
| AMF richness | water × year | 0.024 | 1 | 0.87 |
| | plant richness × year | 0.11 | 1 | 0.74 |
| | plant PDv × year | 0.0051 | 1 | 0.94 |

**Table S2.3**: Model outputs of individual trait correlation with total fungal, AMF, and pathogen richness.

| Trait (all scaled) | Slope | CI_2.5 | CI_97.5 | p.val | p.adj | Community |
|---|---|---|---|---|---|---|
| cwm.LA | -0.061 | -0.120 | -0.002 | 0.041 | 0.48 | Fungi |
| cwm.LMA | 0.007 | -0.051 | 0.065 | 0.811 | 0.95 | Fungi |
| cwm.d15N | 0.037 | -0.021 | 0.094 | 0.209 | 0.68 | Fungi |
| cwm.d13C | 0.005 | -0.053 | 0.062 | 0.873 | 0.95 | Fungi |
| cwm.percN | 0.010 | -0.048 | 0.068 | 0.731 | 0.95 | Fungi |
| cwm.percC | 0.015 | -0.044 | 0.073 | 0.626 | 0.95 | Fungi |
| cwm.CN | -0.010 | -0.067 | 0.048 | 0.742 | 0.95 | Fungi |
| cwm.veg_height | 0.005 | -0.054 | 0.064 | 0.872 | 0.95 | Fungi |
| cwm.rep_height | 0.015 | -0.043 | 0.073 | 0.619 | 0.95 | Fungi |
| fd_dis.r | 0.037 | -0.020 | 0.095 | 0.205 | 0.21 | Fungi |
| cwm.depth | -0.002 | -0.059 | 0.055 | 0.950 | 0.95 | Fungi |
| cwm.avg.diam | -0.043 | -0.100 | 0.013 | 0.133 | 0.58 | Fungi |
| cwm.srl | 0.052 | -0.005 | 0.110 | 0.074 | 0.48 | Fungi |
| cwm.rtd | 0.005 | -0.053 | 0.063 | 0.863 | 0.95 | Fungi |
| fd_dis.s | 0.055 | -0.002 | 0.112 | 0.057 | 0.11 | Fungi |
| cwm.LA | -0.686 | -0.968 | -0.405 | <0.0001 | <0.0001 | AMF |
| cwm.LMA | 0.126 | -0.172 | 0.425 | 0.407 | 0.48 | AMF |
| cwm.d15N | 0.358 | 0.063 | 0.653 | 0.017 | 0.038 | AMF |
| cwm.d13C | 0.146 | -0.154 | 0.446 | 0.341 | 0.48 | AMF |
| cwm.percN | -0.104 | -0.401 | 0.193 | 0.492 | 0.53 | AMF |
| cwm.percC | 0.237 | -0.057 | 0.531 | 0.114 | 0.19 | AMF |
| cwm.CN | 0.137 | -0.161 | 0.435 | 0.367 | 0.48 | AMF |
| cwm.veg_height | 0.301 | 0.010 | 0.592 | 0.043 | 0.079 | AMF |
| cwm.rep_height | 0.500 | 0.212 | 0.788 | 0.001 | 0.002 | AMF |
| fd_dis.r | 0.300 | 0.007 | 0.593 | 0.045 | 0.045 | AMF |
| cwm.depth | -0.051 | -0.348 | 0.246 | 0.737 | 0.74 | AMF |
| cwm.avg.diam | -0.624 | -0.905 | -0.343 | <0.0001 | <0.0001 | AMF |
| cwm.srl | 0.765 | 0.488 | 1.041 | <0.0001 | <0.0001 | AMF |
| cwm.rtd | -0.368 | -0.657 | -0.080 | 0.012 | 0.032 | AMF |
| fd_dis.s | 0.377 | 0.085 | 0.668 | 0.011 | 0.023 | AMF |
| cwm.LA | -0.952 | -3.380 | 1.476 | 0.442 | 0.72 | Pathogen |
| cwm.LMA | -2.079 | -4.491 | 0.332 | 0.091 | 0.29 | Pathogen |
| cwm.d15N | 1.978 | -0.468 | 4.423 | 0.113 | 0.29 | Pathogen |
| cwm.d13C | -0.469 | -2.925 | 1.988 | 0.708 | 0.92 | Pathogen |
| cwm.percN | 2.362 | -0.042 | 4.766 | 0.054 | 0.29 | Pathogen |
| cwm.percC | 0.545 | -1.913 | 3.003 | 0.664 | 0.92 | Pathogen |
| cwm.CN | -2.074 | -4.483 | 0.334 | 0.091 | 0.29 | Pathogen |
| cwm.veg_height | -0.108 | -2.553 | 2.337 | 0.931 | 1.00 | Pathogen |
| cwm.rep_height | -0.006 | -2.450 | 2.438 | 0.996 | 1.00 | Pathogen |
| fd_dis.r | 2.043 | -0.387 | 4.474 | 0.099 | 0.099 | Pathogen |

| | | | | | | |
|---|---|---|---|---|---|---|
| cwm.depth | -1.120 | -3.552 | 1.312 | 0.367 | 0.68 | Pathogen |
| cwm.avg.diam | -2.226 | -4.629 | 0.177 | 0.069 | 0.29 | Pathogen |
| cwm.srl | 1.820 | -0.598 | 4.239 | 0.140 | 0.30 | Pathogen |
| cwm.rtd | -0.154 | -2.588 | 2.280 | 0.901 | 1.00 | Pathogen |
| fd_dis.s | 3.453 | 1.054 | 5.852 | 0.005 | 0.010 | Pathogen |

**Table S2.4**: Interaction between plant diversity (richness or PDv) and trait CWMs on AMF and pathogen richness. We focus on the CWM of traits spanning the fungal collaboration gradient for AMF richness, and those spanning the fast-slow gradient for pathogen richness. None of the interactions were significant.

| Response Variable | Interaction Tested | $\chi^2$ | df | p.val |
|---|---|---|---|---|
| AMF richness | cwm.diam x plant richness | 0.57 | 1 | 0.45 |
| | cwm.diam x plant PDv | 0.77 | 1 | 0.38 |
| | cwm.srl x plant richness | 3.19 | 1 | 0.74 |
| | cwm.srl x plant PDv | 3.35 | 1 | 0.67 |
| Pathogen richness | cwm.rtd x plant richness | 3.66 | 1 | 0.06 |
| | cwm.rtd x plant PDv | 2.25 | 1 | 0.13 |
| | cwm.LA x plant richness | 0.02 | 1 | 0.88 |
| | cwm.LA x plant PDv | 0.21 | 1 | 0.64 |
| | cwm.LMA x plant richness | 0.14 | 1 | 0.70 |
| | cwm.LMA x plant PDv | 0.02 | 1 | 0.88 |
| | cwm.percN x plant richness | 1.23 | 1 | 0.27 |
| | cwm.percN x plant PDv | 0.17 | 1 | 0.68 |
| | cwm.percC x plant richness | 0.10 | 1 | 0.76 |
| | cwm.percC x plant PDv | 0.49 | 1 | 0.48 |
| | cwm.CN x plant richness | 1.10 | 1 | 0.29 |
| | cwm.CN x plant PDv | 0.09 | 1 | 0.76 |

**Figure S2.1**: Remaking Figure 3 when only considering probable and highly probable (excluding possible) plant pathogens in soil. Pathogen richness was no longer correlated with CWMs of any individual traits.

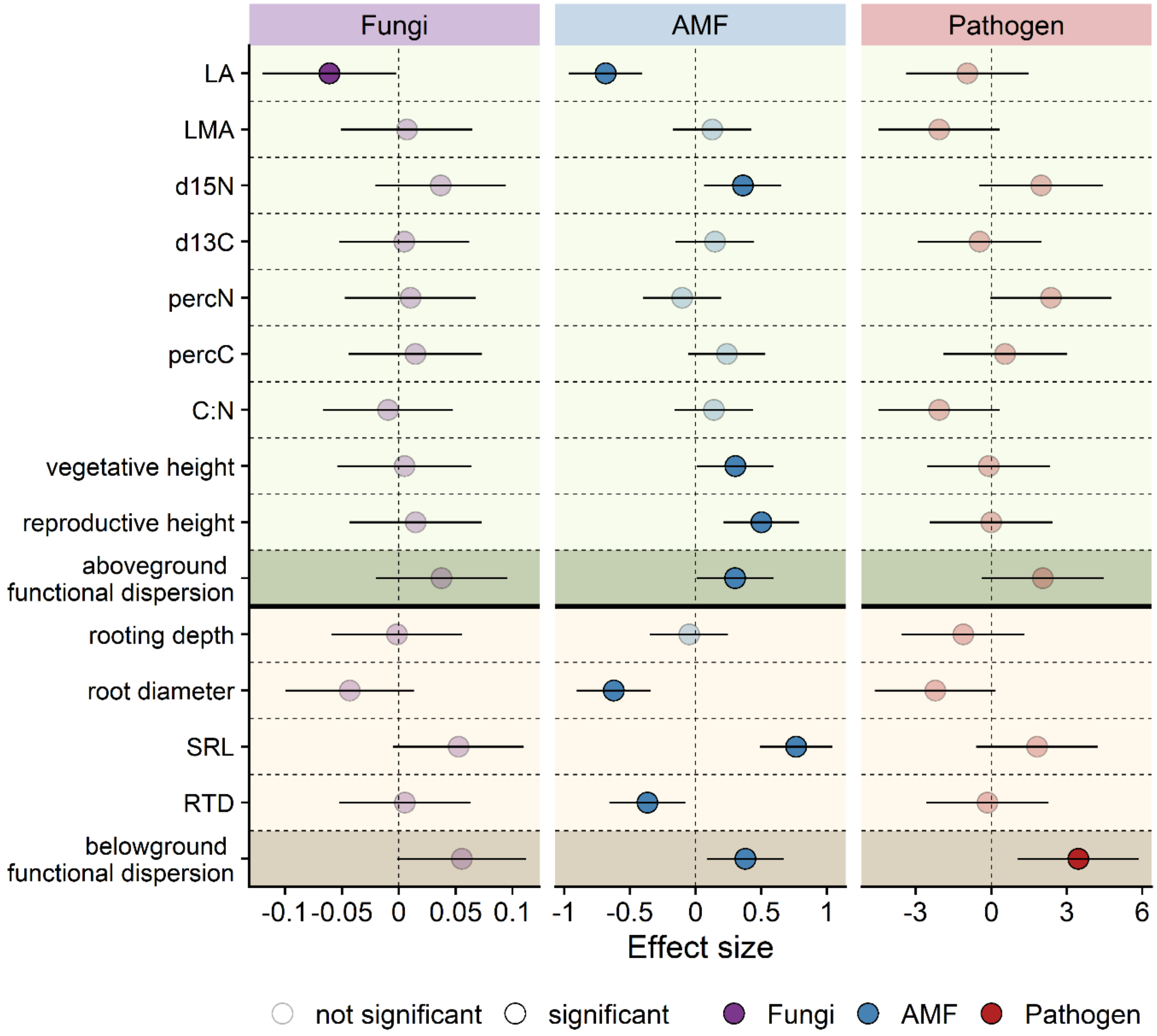

**Figure S2.2**: Remaking Figure 3 after adjusting p values using the Benjamini-Hochberg procedure. Pathogen richness was no longer correlated with CWMs of any individual traits.

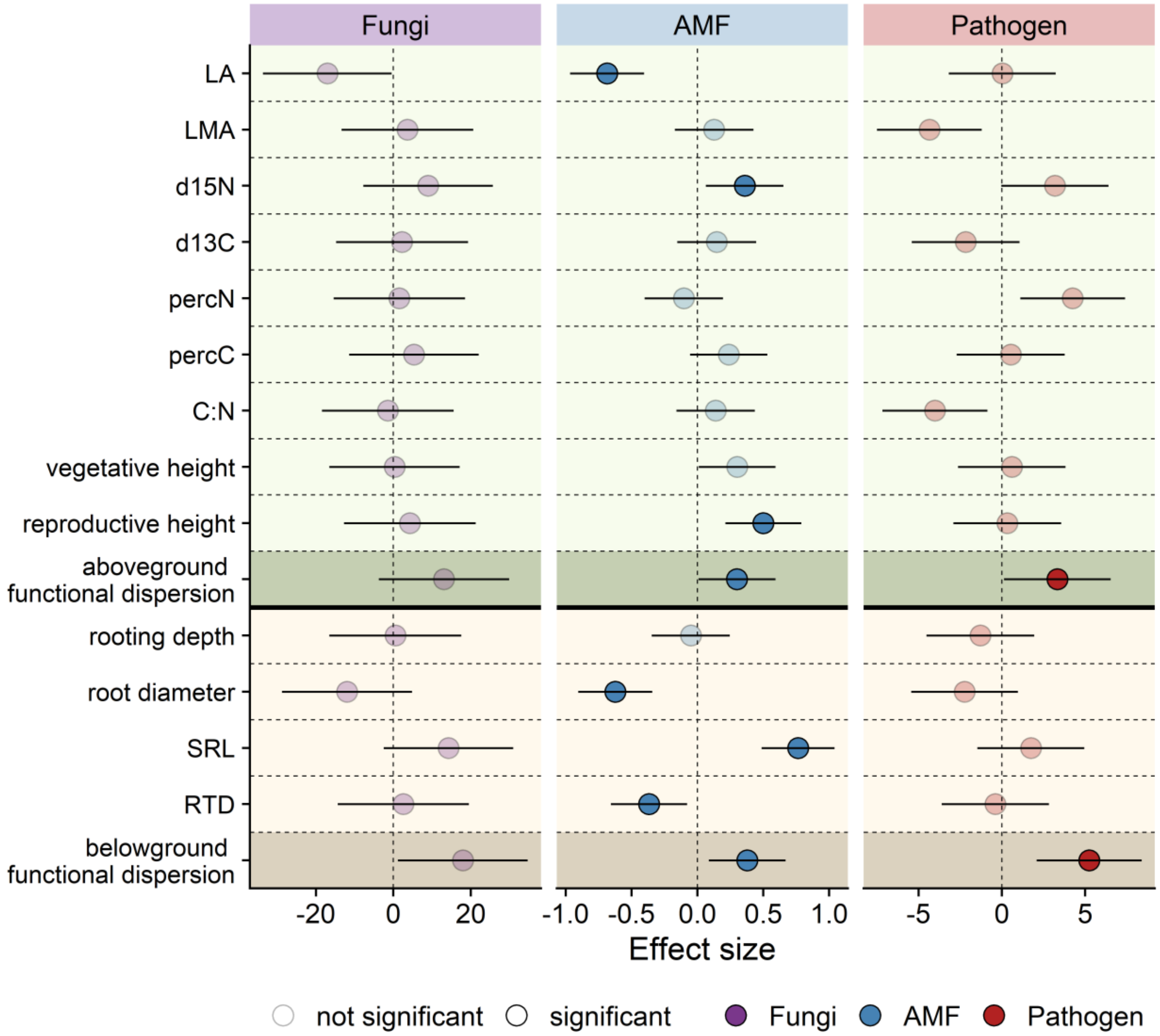

**Figure S2.3**: Remaking Figure 4D with the inclusion of an outlier. The outlier made the significant relationship between AMF richness and plant community complementarity no longer significant

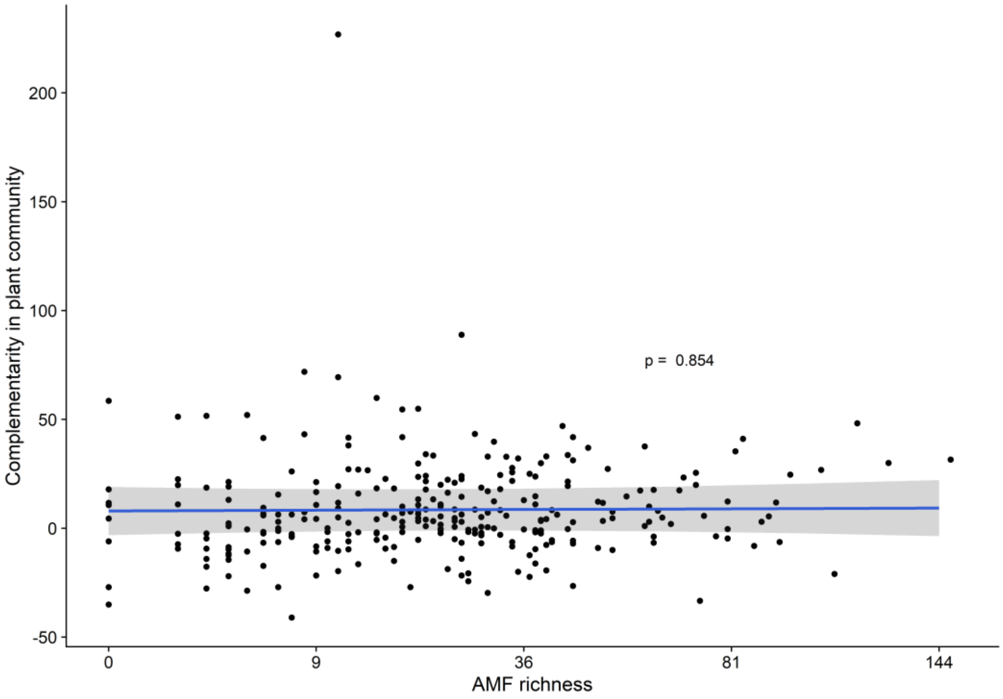
